# High temperature cycles result in maternal transmission and dengue infection differences between *Wolbachia* strains in *Aedes aegypti*

**DOI:** 10.1101/2020.11.25.397604

**Authors:** Maria Vittoria Mancini, Thomas H. Ant, Christie S. Herd, Daniel D. Gingell, Shivan M. Murdochy, Enock Mararo, Steven P. Sinkins

## Abstract

Environmental factors play a crucial role in the population dynamics of arthropod endosymbionts, and therefore in the deployment of *Wolbachia* symbionts for the control of dengue arboviruses. The potential of *Wolbachia* to invade, persist and block virus transmission depends in part on its intracellular density. Several recent studies have highlighted the importance of larval rearing temperature in modulating *Wolbachia* densities in adults, suggesting that elevated temperatures can severely impact some strains, while having little effect on others. The effect of a replicated tropical heat cycle on *Wolbachia* density and levels of virus blocking was assessed using *Aedes aegypti* lines carrying strains *w*Mel and *w*AlbB, two *Wolbachia* strains currently used for dengue control. Impacts on intracellular density, maternal transmission fidelity and dengue inhibition capacity were observed for *w*Mel. In contrast *w*AlbB-carrying *Ae. aegypti* maintained a relatively constant intracellular density at high temperatures and conserved its capacity to inhibit dengue. Following larval heat treatment, *w*Mel showed a degree of density recovery in aging adults, although this was compromised by elevated air temperatures. When choosing the *Wolbachia* strain to be used in a dengue control programme it is important to consider the effects of environmental temperatures on invasiveness and virus inhibition.

**Author summary:** In the past decades, dengue incidence has dramatically increased all over the world. An emerging dengue control strategy utilizes *Ae. aegypti* mosquitoes artificially transinfected with the bacterial symbiont *Wolbachia*, with the ultimate aim of replacing wild mosquito populations. *Wolbachia* is transmitted from mother to offspring and is able to interfere with virus transmission within the mosquito vector. However, the rearing temperature of mosquito larvae is known to impact on some *Wolbachia* strains. In this study, we compared the effects of a temperature cycle mimicking natural breeding sites in tropical climates on two *Wolbachia* strains, currently used for open field trials. We observed that the strain *w*Mel was susceptible to high larval rearing temperatures, while the strain *w*AlbB resulted to be more stable. These results underlines the importance of understanding the impact of environmental factors on released mosquitoes, in order to ensure the most efficient strategy for dengue control.

## Introduction

*Wolbachia* comprises a diverse genus of maternally inherited bacterial endosymbionts that naturally infect arthropod species, but the major arbovirus mosquito vector *Aedes aegypti* is not a native *Wolbachia* host (1). *Wolbachia* facilitates its spread through host populations by increasing the relative fitness of carriers in various ways, including reproductive manipulations such as cytoplasmic incompatibility (CI). CI occurs when a *Wolbachia*-carrying male mates with a *Wolbachia*-free female, and results in reduced egg hatching. However, artificial transfers have been carried out in the laboratory with a range of *Wolbachia* strains, some of which induce strong CI and greatly reduce the competence of *Ae. aegypti* to transmit arboviruses, including Zika and dengue (2-7). An emerging dengue control strategy utilises CI to spread *Wolbachia* through wild mosquito populations, thereby reducing virus transmission. An increasing number of dengue-endemic countries are incorporating releases of *Wolbachia*-carrying *Ae. aegypti* as part of ongoing dengue control efforts. Open-field release programmes are currently underway in Indonesia, Vietnam, Australia and Malaysia, Colombia and Brazil, with significant reductions in dengue incidence reported (8-10). Several *Wolbachia* strains have been stably introduced into *Ae. aegypti*, with different strains generating distinct fitness and pathogen blocking profiles. In particular, the *Wolbachia* strains *w*AlbB and *w*Mel, native to *Aedes albopictus* and *Drosophila melanogaster*, respectively, display promising characteristics in laboratory studies (2, 5, 7, 11, 12), and both are currently being deployed for dengue control. *w*Mel belongs to the supergroup A *Wolbachia* clade. It provides protection from RNA viruses in its native host (13, 14), and blocks the transmission of dengue (DENV), chikungunya (CHIKV) and Zika (ZIKV) viruses in *Ae. aegypti* (2, 6, 15). The *w*Mel infection has been successfully established in *Ae. aegypti* populations in the cities of Cairns and Townsville in northern Australia, and in Yogyakarta, Indonesia, with data indicating reductions in cases of locally acquired dengue (9, 10, 16, 17). *w*AlbB belongs to the supergroup B *Wolbachia* clade, and also efficiently blocks DENV and ZIKV transmission in *Ae. aegypti* (3, 7, 18). Open-field releases of *w*AlbB in Kuala Lumpur, Malaysia, have resulted in high population frequencies and significant reductions in dengue incidence (8).

The magnitude of *Wolbachia*-mediated virus blocking shows a strong positive correlation with intracellular density (19, 20). Fitness costs also correlate with density, and the high density *w*Au and *w*MelPop strains cause both high fitness costs and strong viral inhibition (2, 7, 21, 22), although there are some exceptions (7, 23). *w*Mel and *w*AlbB reach comparable densities in female *Ae. aegypti*, and under standard laboratory conditions they show approximately equivalent levels of dengue (7, 11) and Zika blocking (7), and both have minimal effects on host fitness (2, 7, 24).

Invasiveness and stability of a *Wolbachia* strain depends primarily on CI induction capacity, maternal transmission efficiency and effects on host fitness. The likelihood that a female will mate with a *Wolbachia*-carrying male and incur the fitness cost of CI increases with *Wolbachia* frequency. The fitness advantage of CI is therefore frequency dependent, with invasiveness following bi-stable dynamics determined by an invasion threshold (25, 26). Above the threshold the fitness advantages of CI overcome other fitness costs and *Wolbachia* will tend to spread; below the threshold fitness costs dominate and *Wolbachia* will tend to be lost. The high density *w*MelPop strain induces strong CI, but results in fitness costs over a range of life history traits, including reductions in longevity and the survival of eggs following periods of desiccated quiescence. *w*MelPop carrying *Ae. aegypti* were released in field sites in Australia and Vietnam and despite reaching high initial infection frequencies, the strain was eventually lost once releases ceased (27).

Exposure of host insects to thermal stress is known to unbalance and perturb long-term symbiotic interactions and their phenotypes (28), and *Wolbachia* frequency in insect populations can fluctuate seasonally and between geographical locations (29, 30). In mosquitoes, several recent studies have demonstrated an impact of larval rearing temperatures on *Wolbachia* density in the resulting adults, with results suggesting that elevated temperatures can significantly reduce the density of some strains (7, 31, 32). *w*Mel appears to be particularly sensitive to high temperatures, with density dropping by several orders of magnitude when larvae are exposed to diurnal heat cycling between 27-37°C. In these experiments, a reduced capacity of male carriers to induce CI and a lower level of maternal transmission were observed, with eventual loss of the strain when high rearing temperatures were maintained for more than one generation (31). In contrast, *w*AlbB was found to be more stable at high temperatures, with little (7) or no (31) reduction in density.

Previous studies have investigated the effects of high larval rearing temperatures on *Wolbachia* density in whole mosquitoes, and have examined effects on the transmission fidelity (7, 31, 32). However, reduced densities also suggest the potential for reduced virus blocking. Here, results are presented from a series of experiments examining the effects of simulated tropical temperatures on *Wolbachia* transmission and dengue blocking. Findings indicate that the *w*Mel strain has both reduced maternal transmission and virus blocking capacity following larval rearing at high temperatures.

## Results

### *Effects of field-simulated temperature cycles on* Wolbachia *density*

Detailed temperature recordings from tropical *Ae. aegypti* larval breeding sites were obtained from a previously published study (33), and a replica-cycle (temp min: 28°C; max: 36°C, **Suppl. Fig. 1**) was generated in the laboratory using a programmable dynamic-temperature cabinet. Larvae from *w*Mel- and *w*AlbB-carrying *Ae. aegypti* lines were reared under either simulated field temperatures, or control conditions (constant 27°C). On eclosion, adult mosquitoes from both treatments were maintained at a constant 27°C. 5-day-old adults were sacrificed, and *Wolbachia* densities assessed (**Fig. 1**). A subset of females were blood-fed, and the resulting progeny exposed to a second round of larval heat treatment, with *Wolbachia* densities in G1 adults assessed 5-days after emergence. Consistent with previous studies (7, 31, 32), the *w*Mel strain was particularly susceptible to density reductions resulting from high temperature rearing, with a significant drop in density from 12.65 ± 5.9 *Wolbachia* per cell (mean ± SD), to 1.4 ± 0.92 *Wolbachia* per cell (*p*<0.001, Mann-Whitney test) after one generation of heat treatment. The *w*AlbB strain maintained a relatively constant density over both generations of heat-treatment (*p* >0.57 for both generations, Mann-Whitney test).

**Figure 1.**
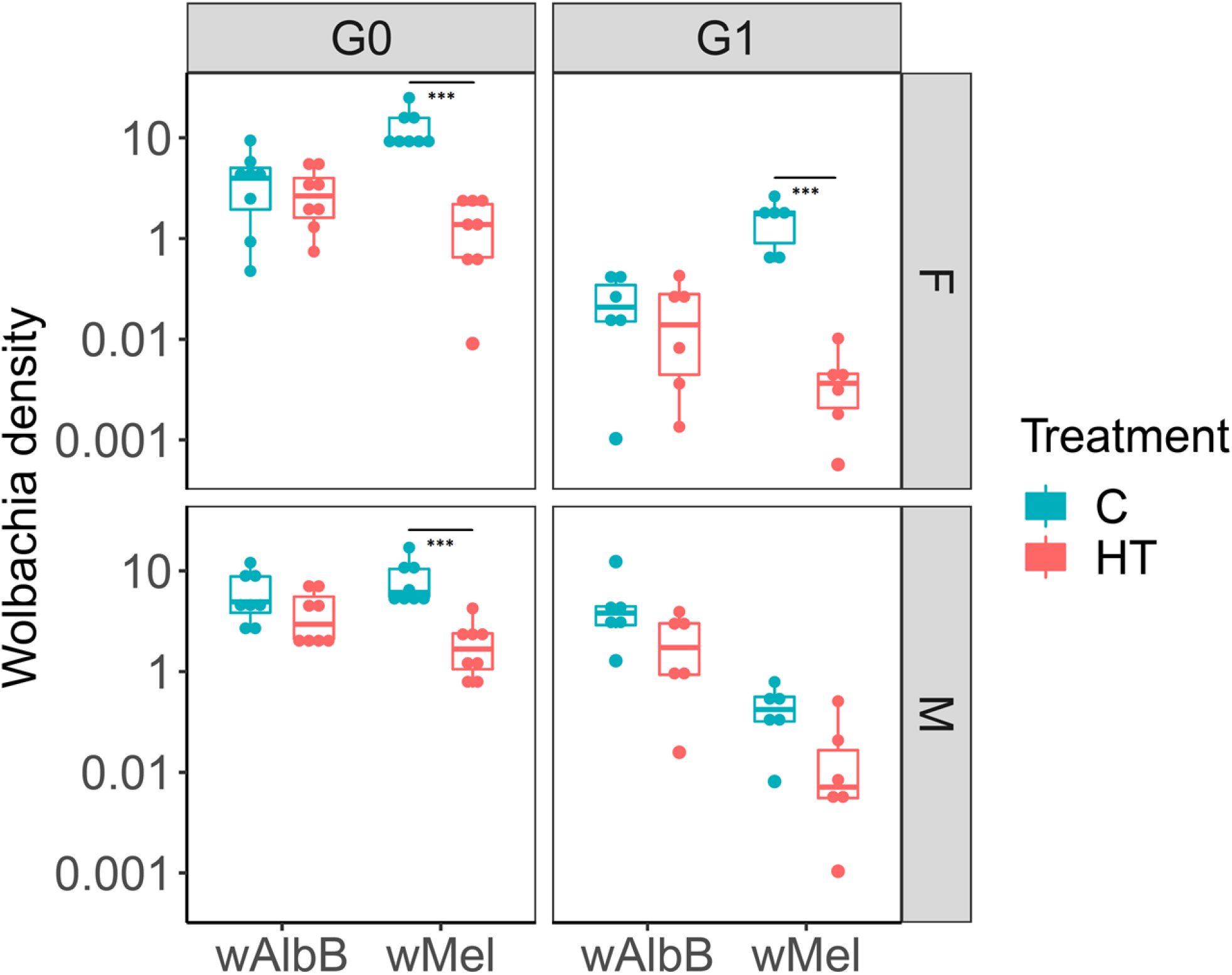
*Wolbachia* density in whole-bodies of control (C, constant 27°C) and heat-treated (HT, temp min= 28°C; temp max= 36°C). The densities of *w*AlbB and *w*Mel were quantified by qPCR on 5-days-old females (F) and males (M) over two generations of heat-treatment. Boxplots represent 6 biological replicates. Central line indicates the median of densities and whiskers represent upper and lower extremes. A Mann-Whitney test was used for statistical analyses.

### *Effects of field-simulated temperature cycles on* Wolbachia *maternal transmission*

*Wolbachia*-carrying females reared under either the tropical high temperature larval temperature cycle or control conditions were back-crossed to wild-type males. Females were individualized for oviposition and the resulting G1 eggs hatched as single families. G1 larvae were reared at a constant 27°C until the L4 stage, whereupon a random selection from each family was assessed for *Wolbachia*-infection status and density. *w*AlbB-females resulting from larvae reared under either tropical high temperature or control conditions transmitted *Wolbachia* to 100% of offspring (N=60). Interestingly, the G1 progeny from heat-treated *w*AlbB mothers tended to show higher *Wolbachia* densities compared to progeny resulting from mothers reared under control conditions; if the densities of larvae from each family are combined, the increase is significant (p<0.001, Mann-Whitney test) (**Fig. 2A**). In contrast, maternal transmission of *w*Mel was significantly reduced following larval heat treatment, with the complete loss of *w*Mel in 3 of the 6 heat-treated families, compared to 100% transmission in the control group (p<0.0001, Fisher’s exact test) (**Fig. 2B**). *w*Mel densities in the *Wolbachia* positive G1 progeny were significantly lower following heat treatment than densities in the G1 progeny following control treatment (*p* =0.002, Mann-Whitney test).

**Figure 2.**
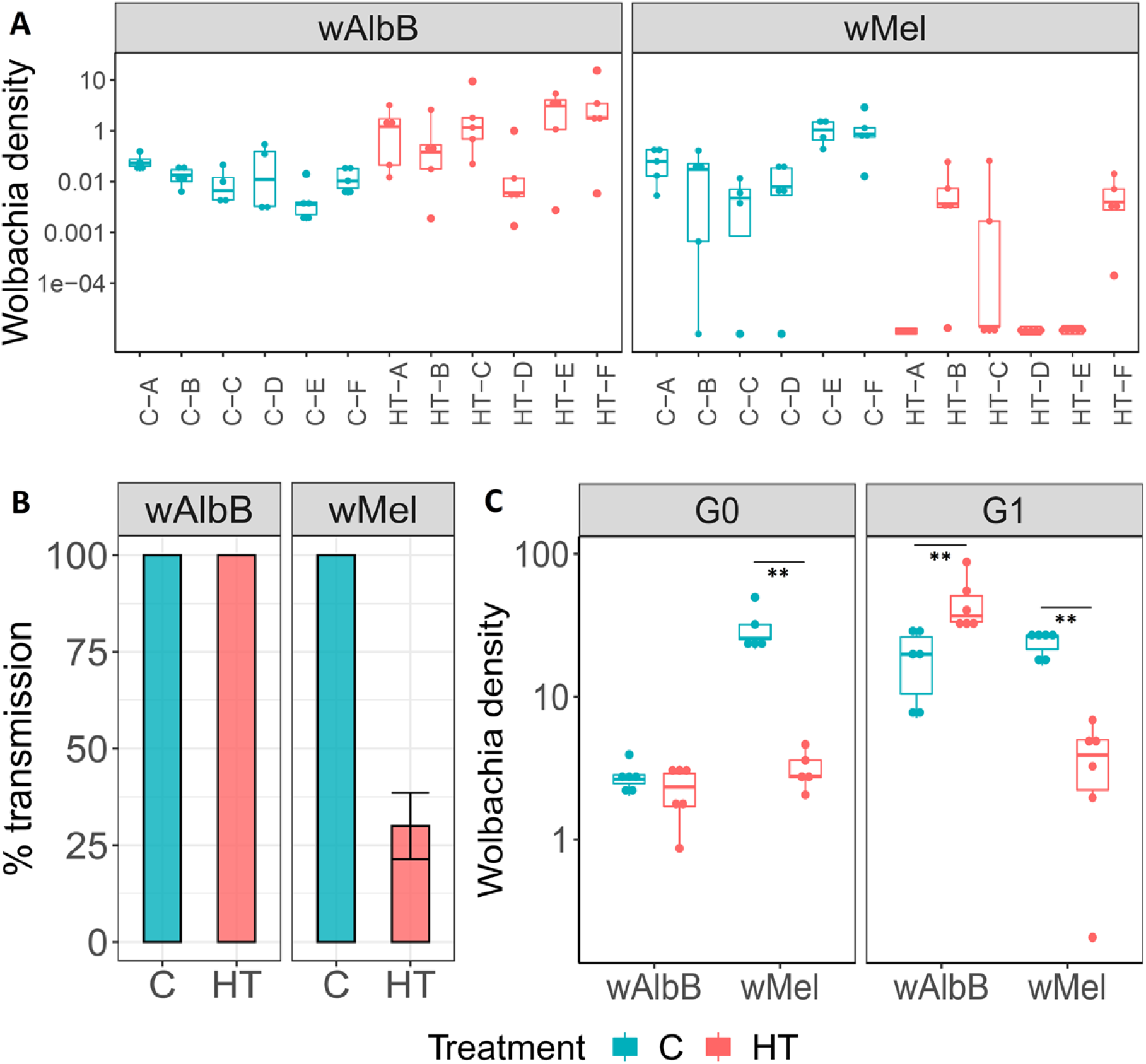
Whole-body densities, maternal transmission rate and ovary-specific densities of *w*AlbB and *w*Mel in control (C, constant 27°C) and heat-treated (HT, temp min=28°C; temp max 36°C) mosquitoes. (**A**) Progeny from single females reared as larvae under control or high temperature conditions were hatched in families and reared at 27°C. 6 L4 larvae were randomly sampled from each individualized female and assessed for *Wolbachia* density by qPCR (**A**) and infection-status by strain-specific PCR (**B**) (N=60 for each treatment/strain). (**C**) Densities of *w*AlbB and *w*Mel were measured in 6 pools of 3 sets of dissected ovaries. The centre of the box-plots indicates the median of densities and whiskers represent upper and lower extremes. A Mann-Whitney test was used for statistical analyses.

To correlate reductions in maternal transmission with *Wolbachia* densities in ovaries, females reared at either high or control larval temperatures were dissected, and ovary densities assessed by qPCR. *Wolbachia* was also visualized in ovaries by whole-mount fluorescent *in situ* hybridisation (FISH) (**Suppl. Fig. 2**). Results indicate that the high temperature-cycle caused significant reductions in the ovary density of *w*Mel, while the density of *w*AlbB was not negatively affected by the high temperature cycle, and even increased (p=0.002, Mann Whitney Test), compared to controls following two generations of treatment (**Fig. 2C**).

### Effects of field-simulated temperature cycles on virus transmission

To test whether temperature-induced reductions in *Wolbachia* density could impact dengue inhibition, larvae from the *w*AlbB, *w*Mel and wild-type lines were reared under either high-temperature or control conditions, and the resulting adult females were orally challenged with a bloodmeal containing DENV-2. 12 days post-feeding, levels of infectious virus in dissected heads and thoraxes, and salivary glands, were quantified to assess the infection rate and transmission potential within the vector.

Increasing the larval rearing temperature had no significant effect on the infection rate in head and thoraxes of wild-type females - 8 out of 24 for the control group and 14 out of 24 for the heat treated cohort (*p=*0.14, Fisher’s exact Test). A slight, although non-significant, increase in viral titres was observed between the groups (*p*=0.05, Mann-Whitney Test). In contrast, *w*Mel females reared at high temperature displayed a significant increase in infection rate in heads and thoraxes (**Fig. 3D**, 8 out of 24 were positive for virus, 33.3%) compared to *w*Mel females reared under control conditions (**Fig. 3B**, 1 out of 24 were positive for virus, 4.2%) (*p*<0.05; Fisher’s Exact Test). While *w*Mel females reared under control conditions had a significantly reduced infection rate (*p*=0.02, Fisher’s Exact Test) compared to wild-type females, this decrease was not observed when *w*Mel larvae were reared at high temperature (*p*=0.14, Fisher’s Exact Test). However, *w*Mel caused a significant reduction in viral titre in heads and thoraxes compared to wild-type *Ae. aegypti*, regardless of larval rearing temperature (*p*= 0.003 for control and *p=*0.03 for heat treated *w*Mel, respectively; Mann-Whitney Test), although viral titres were significantly higher in *w*Mel females when larvae were reared at high-compared with control temperature (*p*=0.01, Mann-Whitney) (**Fig. 3A and C**). Moreover, no significant difference in the viral titer in salivary gland tissue was observed in heat-treated *w*Mel compared to wild-type females (*p*=0.27, Mann-Whitney Test) (**Fig. 3E**).

**Figure 3.**
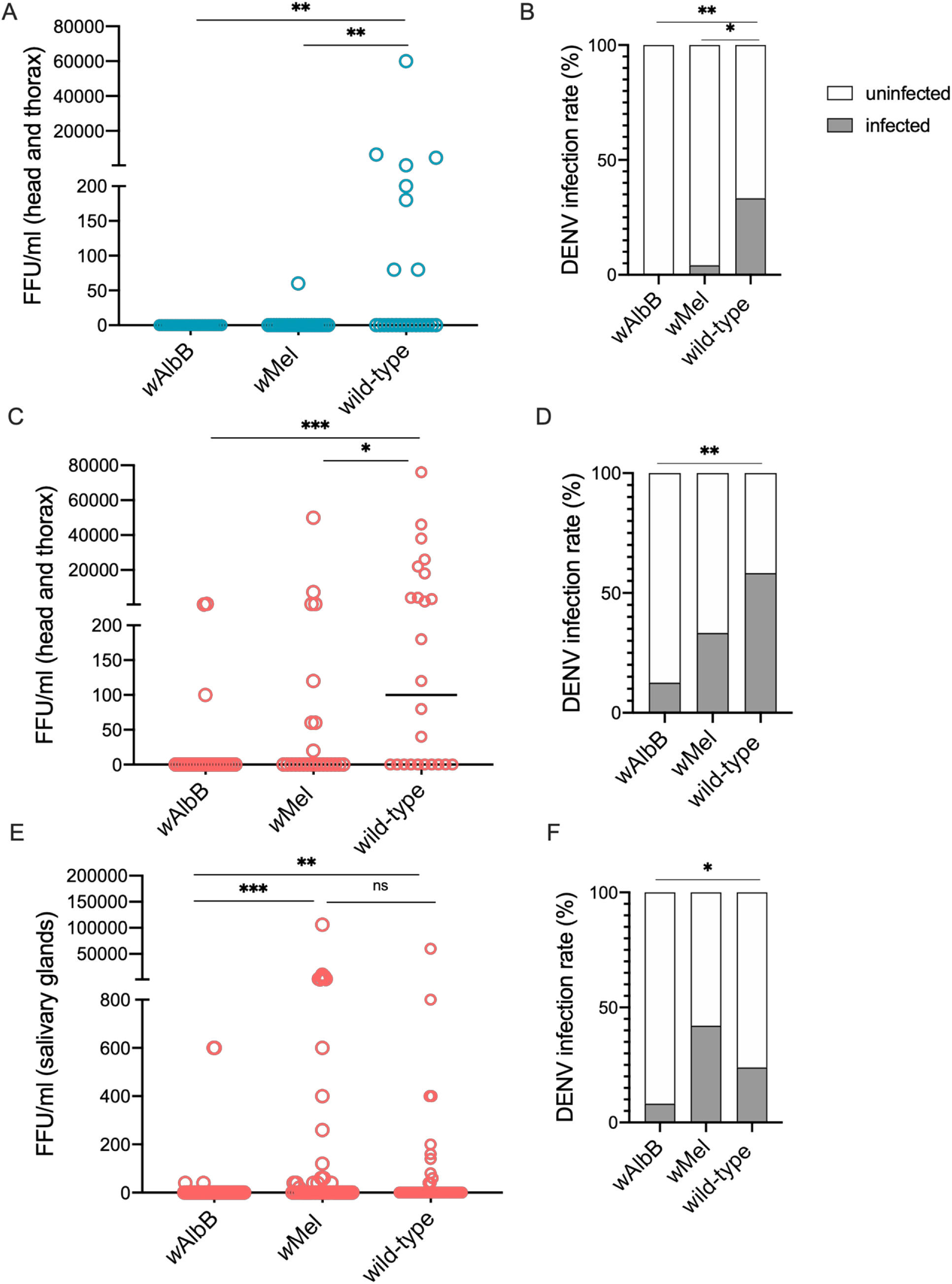
Effect of larval heat-treatment on dengue inhibition. Wild-type, *w*AlbB- and *w*Mel-carrying females were fed on DENV2-infected blood-meal. Engorged females were selected and incubated for 12 days. Heads and thoraxes from control (**A**) and heat-treated females (**C)** were assessed for virus dissemination by Fluorescent Focus Assay (FFA). Viral titer was also assessed on salivary glands of heat-treated females from an independent viral challenge (**E**). Dots represent the number of foci/ml and each dot corresponds to a single mosquito. Dissemination rates from the same experiments are represented in panels **B, D** and **F**. Statistical analysis was performed using Mann-Whitey Test and Fisher’s Exact Test. See Table S2 for statistical comparisons.

**Figure 4.**
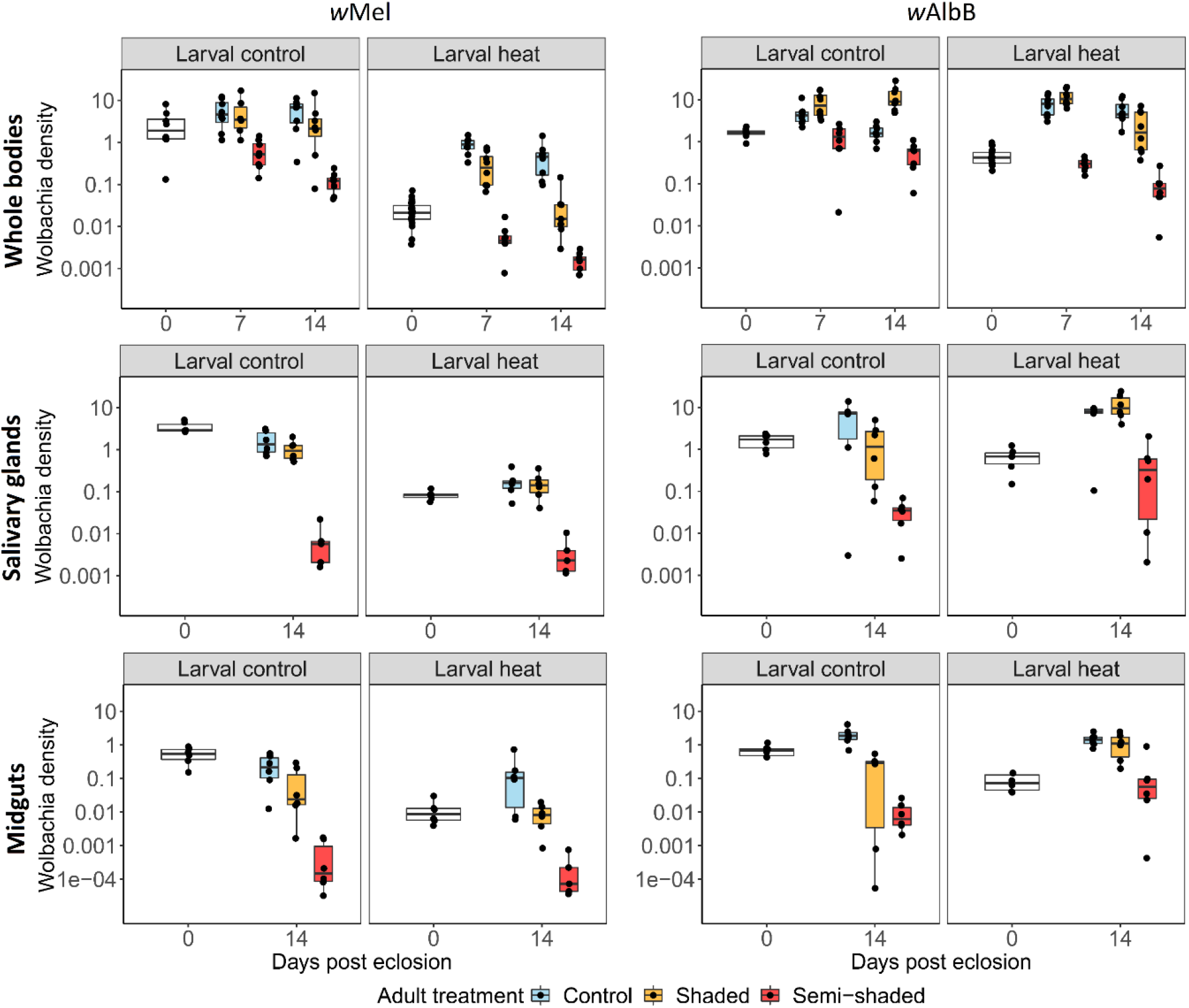
Effects of high temperature larval and adult ambient air temperatures on *w*Mel and *w*AlbB densities in whole-bodies, salivary-gland and midgut tissues. Larvae were reared under control (larval control, constant 27°C) and high temperature (larval heat, temp min= 27°C; temp max 37°C) conditions. A subset of females were sampled immediately on eclosion (day 0), and densities assessed. The remaining females were divided into three adult treatment temperatures: control (constant 27°C), shaded (temp min = 28°C; temp max = 33.5°C), and semi-shaded (temp min = 27°C; temp max = 36.5°C). Adults were sampled and densities assessed in whole bodies (days 7 and 14 post eclosion) and dissected salivary gland and midgut tissues (day 14 post eclosion). Data points represent single whole adult females, or pools of three salivary glands or midguts.

*w*AlbB maintained strong viral inhibition following high-temperature rearing, with no significant difference in DENV-2 infection rate in heads and thoraxes in control-reared (0 out of 24 positive for virus) and high-temperature-reared *w*AlbB females (3 out of 24 positive for virus, 12.5%) (*p*=0.238, Fisher’s Exact Test). Regardless of larval rearing temperature, *w*AlbB consistently reduced the infection rate (control: *p*=0.003, Fisher’s Exact Test; high-temperature: *p*=0.002, Mann-Whitney) and titre of DENV-2 in heads and thoraxes compared to wild-type females (control: *p*=0.003, Fisher’s Exact Test; high-temperature: *p*<0.001, Mann Whitney Test). Moreover, viral titres in salivary gland tissue were significantly lower in high-temperature-reared *w*AlbB females compared to wild-type females (*p*<0.01, Mann-Whitney Test) (**Fig. 3C and E**). Viral infection rate and titre were significantly higher in the salivary glands of heat treated *w*Mel females compared to *w*AlbB-carrying mosquitoes reared at similar high temperatures (p=0.0001, Fisher’s Exact Test; p<0.0001, Mann-Whitney Test), while no significant difference was observed in heads and thoraxes of control and heat treated females from the two *Wolbachia* strains.

### Adult exposure to elevated temperatures and Wolbachia recovery

A previous study reported substantial recovery of *w*Mel in adult *Ae. aegypti* from initially low densities following larval rearing at high temperatures (32). To further investigate density recovery in adults, and to examine the impact of elevated air temperatures on recovery rates, *w*Mel and *w*AlbB larvae were reared under control or high temperature conditions, with emerging adults exposed to replica heat cycles generated from recordings of ambient temperatures in shaded (temp min: 28°C; max: 33.5°C) or semi-shaded (temp min: 27°C; max: 36.5°C) sites in urban Kuala Lumpur (**Suppl. Fig. 1)**. Adult females were dissected, and *Wolbachia* densities in midgut and salivary gland tissues were also assessed.

There was a significant reduction in the density of *w*Mel in adults emerging from the high temperature cycle (0.025 ± 0.015 *Wolbachia* per cell, mean ± SD) compared to larval-control treatments (2.8 ± 2.6 *Wolbachia* per cell) (*p* <0.001, Mann-Whitney Test). However, there was a marked recovery in density in heat-treated larvae subsequently reared under control conditions as adults (reaching 0.495 ± 0.435 *Wolbachia* per cell after 14 days), although this recovery was incomplete – with adults from control larvae maintaining a significantly higher density, 5.975 ± 3.73 *Wolbachia* per cell after 14 days (*p* =0.003, Mann-Whitney Test). Air temperature had a significant impact on the recovery of *w*Mel, with 14-day-old females from the shaded and semi-shaded cycles displaying significantly lower densities than adults reared at control temperatures (*p* <0.001 for both shaded and semi-shaded, Mann-Whitney test).

A similar trend was observed in dissected salivary gland and midgut tissues of emerging adults, with significant reductions in *w*Mel density in both tissues following larval heat treatment (*p* <0.005 for both midguts and salivary glands, Mann-Whitney test). There was a recovery in density in midguts at day-14 in the larval-heat cohort reared at the control adult temperature, with no significant difference compared to mosquitoes reared exclusively under control conditions (*p* =0.48, Mann-Whitney test). Densities in the salivary glands of females reared under larval-heat showed minimal recovery at both control and shaded adult treatments, with significantly lower densities compared to mosquitoes reared exclusively under control conditions (*p* <0.005 for both control and shaded, Mann-Whitney test). Adults reared under the semi-shaded heat cycle showed significant reductions in density in both salivary gland and midgut tissues compared to adults reared at control temperatures (*p* <0.005 for both salivary glands and midguts, Mann-Whitney test).

*w*AlbB showed a reduction in density in adults emerging from larval-heat (0.46 ± 0.2 *Wolbachia* per cell) compared to larval-control treatments (1.65 ± 0.41 *Wolbachia* per cell) (*p* <0.001, Mann-Whitney test). However, the density recovered fully after 7-days of adult rearing under control conditions, with no significant reduction in density compared to mosquitoes reared only at control temperatures (*p*=0.002, Mann-Whitney test). Interestingly, *w*AlbB-carriers reared at the control temperature as larvae and the shaded temperature cycle as adults showed significantly increased *Wolbachia* densities compared to adults reared only under control conditions (*p* =0.002, Mann-Whitney test). The semi-shaded adult treatment resulted in significant reductions in densities compared to adults reared at the control temperature, regardless of larval treatment (*p* <0.001 for both control and heat-treated, Mann-Whitney test).

Following larval heat treatment, the density of *w*AlbB in both midgut and salivary gland tissues of eclosing adults was slightly but significantly reduced (*p* =0.02 for salivary glands; *p* =0.002 for midguts). However, densities in both tissues recovered fully at day-14 when adults were reared under either control or shaded temperature conditions, displaying no significant reductions compared to tissue densities in adults reared under control conditions only (*p* >0.18 for all comparisons). Similar to *w*Mel, rearing *w*AlbB adults under semi-shaded conditions resulted in significant reductions in densities compared to adults reared under control conditions, regardless of larval treatment.

## Discussion

*Ae. aegypti* larvae developing in the tropics encounter a far broader and variable range of temperatures than those typically used in mosquito insectaries (usually stringently maintained in the range of 27-28°C). Several recent studies have highlighted the substantial influence that larval water temperature has in determining the density of some *Wolbachia* strains in *Ae. aegypti*, particularly *w*Mel (31, 32, 34). This is noteworthy as *Wolbachia* strain characterisation is routinely performed under standard insectary temperatures and suggests that phenotypes predicted by laboratory tests may vary in the field. Tropical breeding sites can experience heating above 30°C for extended periods of the day, and in some cases reach daily maxima in excess of 36°C (33, 35). The high temperature regime used in this study was generated from data collected from water drums in Trinidad acting as *Ae. aegypti* larval habitats (33).

Consistent with previous studies, *w*Mel was found to be negatively affected by exposure to the high temperature cycle, showing a significant decrease in whole body density. A substantial drop in adult ovary density was also observed, leading to a reduction in maternal transmission of approximately 75%. Imperfect maternal transmission can impact the population stability of a *Wolbachia* infection by increasing invasive threshold, potentially compromising the ability of *w*Mel to spread and persist in wild populations. Previous evidence documented a disruption in *w*Mel maternal transmission and CI induction following exposure to high temperatures (31). Additionally, intense artificial laboratory selection for a heat resistant *w*Mel variant in *Ae. aegypti* failed to produce a strain with improved thermal tolerance, an observation that was supported by experiments showing that field collected *w*Mel-carriers from a hot climate did not differ substantially in their response to heat stress compared to a laboratory colony – suggesting that adaption of the strain to high temperatures may be intrinsically difficult (34). In contrast, the *w*AlbB strain showed relative heat stability when larvae were reared under the high temperature regime. High densities were maintained in the ovaries, resulting in complete maternal transmission, suggesting that the *w*AlbB strain would be more stable in hot tropical climates.

For the first time, the consequences of tropical heat stress on the ability of *Wolbachia* to inhibit dengue virus dissemination was tested in *w*Mel and *w*AlbB-carrying *Ae. aegypti*. Rates of mosquito infection to salivary glands following challenge with DENV2 were quantified in order to predict the infective state of mosquitoes reared under either high temperature or control conditions. Following exposure to thermal stress, *w*AlbB retained its ability to efficiently block DENV2 dissemination, while *w*Mel showed a significant increase in viral dissemination. *Wolbachia*-mediated viral inhibition is thought to be primarily cell autonomous (5, 36); consequently, densities in midgut and salivary gland tissues are key to blocking virus dissemination and transmission. The reduction in dengue inhibition in heat-treated *w*Mel is concomitant with large reductions in *Wolbachia* density in both midgut and salivary gland tissues, although the density in midguts appeared to recover in adults after 14-days. *w*AlbB also showed reductions in density in midgut and salivary gland tissues, although the reduction was not as dramatic as *w*Mel, and recovered fully in 14-day old adults.

A decrease in the efficiency of dengue blocking by *w*Mel could have significant impacts on the utility of the strain as a vector control intervention in hot tropical climates. This is particularly relevant given the role of high temperatures as a covariate of dengue transmission (37). Moreover, there is the potential that a weakening of the *w*Mel transmission blocking phenotype following exposure to high temperatures could increase the risk of selection of virus escape mutations that confer a lower general susceptibility to *Wolbachia*-mediated inhibition – and could therefore undermine *Wolbachia* interventions. *Wolbachia* at high density induce a broad range of perturbations in *Ae. aegypti* cells (38), including in a number of pathways that are important in the flavivirus life cycle – such as lipid transport and metabolism, autophagy, vesicular trafficking and endoplasmic reticulum stress; this is inherently likely to reduce the risk of selection of virus escape mutations; however, at lower density the levels of perturbation are reduced (38).

A previous study has shown that initially low densities of *w*Mel following larval heat-treatment can recover substantially in adults reared under normal insectary conditions (32). Results presented here are consistent with this finding, with *w*Mel showing considerable (although incomplete) density recovery when adults are reared at a constant 27°C. However, while adult mosquitoes are able to fly and seek cooler resting areas, ambient air temperatures are often very high in the tropics. Recordings in shaded and semi-shaded sites from urban Kuala Lumpur indicate that air temperatures can reach in excess of 34°C for several hours of the day. In larvae carrying *w*Mel reared using the high temperature cycle, and subsequently reared as adults using a replica shaded air-temperature cycle, only a limited recovery in *Wolbachia* density occurred. In contrast, the density of *w*AlbB in whole mosquitoes reared as adults using the shaded temperature cycle were significantly higher than controls – suggesting that the temperature optimal for *w*AlbB replication may actually be higher than the 27°C used in standard rearing. Both *w*Mel and *w*AlbB densities were substantially reduced by exposure to semi-shaded equivalent air temperatures, suggesting that *w*AlbB is not completely resistant to the effects of high temperatures, although this cycle represents an extreme temperature regime that adult mosquitoes will be unlikely to encounter for extended periods. The *w*AlbB strain was capable of reaching and maintaining high frequencies and significantly reducing dengue transmission in the hot tropical climate of urban Kuala Lumpur, Malaysia (8).

Although *w*Mel shows reduced densities in the laboratory using simulated field conditions, releases in Australia, Brazil and Indonesia demonstrate that *w*Mel can stably invade wild *Ae. aegypti* populations (9, 10, 16, 39, 40) and maintain its ability to block dengue (17, 41). In direct comparisons the *w*Mel line produced slightly lower fitness costs than *w*AlbB (7), suggesting that it may be the more invasive of the two strains in cooler climates. Exposure of *w*Mel-carrying *Ae. aegypti* adults to a diurnal temperature cycle with a mean of 28°C and a fluctuating range of 8°C (±4°C) caused a decrease in bacterial density when compared to constant 25°C, but did not reduce the ability of *Wolbachia* to inhibit dengue transmission (42). Moreover, in some hotter equatorial areas *Ae. aegypti* can exploit underground larval habitats, such as wells and drains, which will be away from direct sun light and cooler than ground level. Laboratory experiments have also proved that the effects of thermal stress on *Wolbachia* density are stage-specific (43); in particular, exposure of early larval stage generates a significant and irreversible decrease in density, while the drop observed during exposure to later stages is rescued during adulthood. This suggests that the variations in temperature typical of the field will result in a more complex gradient of phenotypes, less clear-cut than those produced in the laboratory. The complex interactions between environmental temperatures and *Wolbachia* phenotypes has been recently investigated in natural *Wolbachia*-*Drosophila* associations, where the developmental temperature of the host modulated *Wolbachia*-induced antiviral effects, ranging from complete to no protection, although without affecting its density (44).

Our data show that high tropical temperatures have a significant impact on the phenotypic stability of *Wolbachia* in *Ae. aegypti*, and the magnitude of this impact varies substantially between *Wolbachia* strains. Of the strains currently used in open field releases, *w*Mel appears to be particularly susceptible and *w*AlbB relatively stable under thermal stress, with *w*Mel displaying a marked reduction in capacity for maternal transmission and dengue blocking - which is not observed with *w*AlbB. The selection for the optimal strain for *Wolbachia*-deployed vector control strategies must therefore consider phenotypic stability in relation to the geography and climate of selected intervention areas. The water temperature of natural breeding sites not only represents a crucial abiotic factor known to directly affect vector biology (45), but it also plays a role in ensuring the most effective *Wolbachia*-based strategy for reducing dengue transmission.

## Methods

### Mosquito rearing

*w*Mel, *w*AlbB and wild-type *Ae. aegypti* mosquitoes were derived from previously generated lines (7). Colonies were maintained at 27°C and 70% relative humidity with a 12-hour light/dark cycle. Larvae were fed with tropical fish pellets (Tetramin, Tetra, Melle, Germany) and adults maintained with 5% sucrose solution *ad libitum*. Blood meals were provided using an artificial blood-feeding system (Hemotek, UK) using human blood (Scottish National Blood Transfusion Service, UK). Eggs were collected on a wet filter-paper (Grade 1 filter paper, Whatman plc, GE healthcare, UK). Eggs were desiccated for 5 days and later hatched in deionized water containing 1g/L bovine liver powder (MP Biomedicals, Santa Ana, California, USA).

### Temperature cycles

For each replicate, eggs from *w*Mel, *w*AlbB and wild-type (WT) *Ae. aegypti* lines were hatched and separated into experimental groups: larval density (200 larvae per 500 mL of water) and food were consistent between the conditions. Data shown in the plots are the representation of one of three independent biological replicates, consistently showing the same trend of results. Heat-challenged larvae were maintained in Panasonic MLR-352-H Plant Growth Chamber incubator (Panasonic, Osaka, Japan). The applied temperature regime was based on data from *Ae. aegypti* larval breeding containers in Trinidad (33) and replicated in the cabinets. Water temperatures were continuously monitored using a data logger (Hobo Water Temperature Pro V2, Bourne, MA, USA) placed in a plastic tray filled with 500 ml of water. Temperature data were registered and monitored. Mosquitoes under control conditions were stably maintained at 27°C, as previously described.

Pupae were sexed according to size, introduced into cages and maintained during the adult stage at 27°C, unless otherwise stated.

For assessing *Wolbachia* recovery during the adult stage, females from control and heat-treated groups were selected and divided into three different adult treatments: i) C: control (27°C constant), ii) S: shaded (temperature peak at 32°C) and iii) S/S: semi-shaded (temperature peak at 37°C). Adults temperature cycles are based on air temperature readings registered in Kuala Lumpur in February 2019. Readings for the shaded cycle (S) were collected in the area of Pusat Komersial Shah Alam (3°03’57.2”N 101°29’24.0”E), and in the area of the Institute of Medical Research (3°10’10.3”N 101°41’55.0”E) for the semi-shaded cycle (S/S).

### *Wolbachia* density and Fluorescent In Situ Hybridization (FISH)

Genomic DNA from 5-7-days old (unless otherwise stated) whole females and males of *Wolbachia*-carrying lines was extracted with STE buffer (10uM Tris HCL pH 8, 100mm NaCl, 1mm EDTA) and used for *Wolbachia* density quantification by qPCR using the relative quantification of the *Wolbachia* surface protein (*wsp*) gene against the homothorax gene (HTH) as reference gene. The following program was used to run the qPCRs: 95 **°**C for 5 min, 40× cycles of 95 **°**C for 15 s and 60 **°**C for 30 s, followed by a melt-curve analysis. A Rotor Gene Q (Qiagen) was used with 2x QuantiNova SYBR.

Ovaries, salivary glands and midguts (6 pools of 3 organs per each replicate) were dissected from 5-days old females using sterile forceps and needles in a drop of sterile PBS buffer, and immediately transferred into tubes containing STE buffer; genomic DNA from tissues was extracted and *Wolbachia* density was assessed by qPCR as previously described.

At the same time, ovaries were also dissected for Fluorescent *In Situ* Hybridization (FISH) in sterile PBS buffer, and then immediately transferred to a tube containing Carnoy’s fixative (chloroform:ethanol:acetic acid, 6:3:1) and fixed at 4°C overnight. Samples were then rinsed in PBS and incubated in a hybridization buffer containing: 50% formamide, 25% 20xSSC, 0.2% (w/v) Dextran Sulphate, 2.5% Herring Sperm DNA, 1% (w/v) tRNA, 0.015% (w/v) DTT, 1% Denhardt’s solution, and 100 ng/ml of each probe. The probes annealed on the *wsp* gene(5). Samples were left to hybridize overnight in a dark-humid box at 37°C. Samples were washed twice in a solution containing: 5% 20xSSC, 0.015% (w/v) DTT, and twice in a solution of 2.5% SSC, 0.015% (w/v) DTT in dH2O, and incubated at 55°C for 20 minutes. Samples were then placed on a slide containing a drop of VECTASHIELD Antifade Mounting Medium with DAPI (Vector Laboratories, California, USA) and were visualized immediately using a confocal microscope (ZEISS, Germany)

### *Wolbachia* recovery during adult stage

After larval treatments, named control (C) and heat-treated (HT), 8 females from different experimental groups denoted as larval-treatment/adult-treatment as follows: control/control; control/shaded; control/semi-shaded; heat/control; heat/shaded; heat/semi-shaded, were sampled after 0, 7 and 14 days. Midguts and salivary glands were also dissected a few hours after eclosion (day 0) and after 14 days. *Wolbachia* density was assessed in whole mosquitoes and tissues by qPCR as previously described.

### Maternal transmission

Maternal transmission of each *Wolbachia* strain after heat stress was evaluated by backcrossing heat-treated females with heat-treated wild-type males, while control females mated with control wild-type males. After offering a blood-meal, 10 engorged females per group were selected and, after 3 days, individualized on damp circle of filter paper inside up-turned plastic cups. Filter papers were collected and individually desiccated. Once dried, eggs were hatched in containers and reared at stable control temperature; 6-10 4th-instar larvae were randomly sampled from each individualized female (10 females) and assessed for *Wolbachia* infection by PCR, using strain specific primers described in Table S1. PCR reactions were set up using 1x Taqmaster mix (Vazyme) according to the manufacture’s protocol; the amplification reaction consisted of a cycle at 94 °C for 3 min, followed by 30 cycles of denaturation at 94 °C for 30 s, annealing at 55 °C for 30 s, extension at 72 °C for 30s, and a final step at 72 °C for 10 min. Additionally, a subset of samples (6 individuals for 6 family) were validated by qPCR, using *wsp* general primers.

### Virus challenge

5 days-old females per group were fed an infectious blood-meal consisting of human blood and DENV serotype-2 virus (New Guinea C Strain). The virus was serially passaged in *Ae. albopictus* C6/36 cells: the infected supernatant was harvested, concentrated using Amicon Ultra-15 filters (Millipore, IRL) and titered *via* fluorescent focus assay (FFA), as described below. Two independent challenges were carried out, using the same batch of propagated virus at the final concentration in the blood of 1.7x 10^7^ FFU/ml. Control and heat-treated females were infected during the first virus challenge, while the second infectious feeding involved only heat-treated individuals. Fully engorged females were transferred in a climatic chamber at 27°C, 70% relative humidity and a 12-hour light/dark cycle, and maintained with 5% sucrose solution. After 12 days, mosquitoes were dissected and sampled: for the first replicate head thoraxes of control and heat-treated females were used to quantify virus titres, while salivary glands from heat-treated females were used for the second replicate. Samples were transferred in Dulbecco’s Modified Eagle Medium (DMEM) medium supplemented with 2% fetal bovine serum (FBS), After being homogenized, 10-fold serial dilutions (10^−1^ to 10^−3^) of the each solution were transferred onto a monolayer of Vero cells for viral quantification with fluorescent focus assay (FFA). Primary antibody for DENV was MAB8705 Anti-dengue virus complex antibody (Millipore); secondary antibody was the Goat anti-mouse Alexa Fluor 488, A-11001 (Thermo Scientific, Waltham, Massachusetts, USA). Celigo Imaging Cytometer (Nexcelom Bioscience, Lawrence, Massachusetts) was used for imaging plates. Fluorescent foci were counted by eye (from dilutions with less than 100 foci) and virus titers calculated and expressed as FFU/mL.

### Statistical analysis

Graphics were generated using the ‘ggplot2’ package of R Studio (RStudio Inc., Boston, Massachusetts, USA) of the R software (version 3.6.1) and Prism Software (version 8.4.3). All statistical analyses were run using Prism version 8. Shapiro-Wilk Test was used for assessing normality distribution of data, and parametric and non-parametric tests were selected accordingly. Analysis of virus-challenged mosquitoes was performed using a non-parametric Mann-Whitney Test between viral titres and Fisher’s Exact Test for comparing rates of positive and negative samples.

## Acknowledgments

We thank Ghazali M. R. Kamarul, Institute for Medical Research, Malaysia, for providing the temperature readings from Kuala Lumpur and Ary Hoffmann, University of Melbourne for comments on the manuscript.

---

**S1 Figure.**
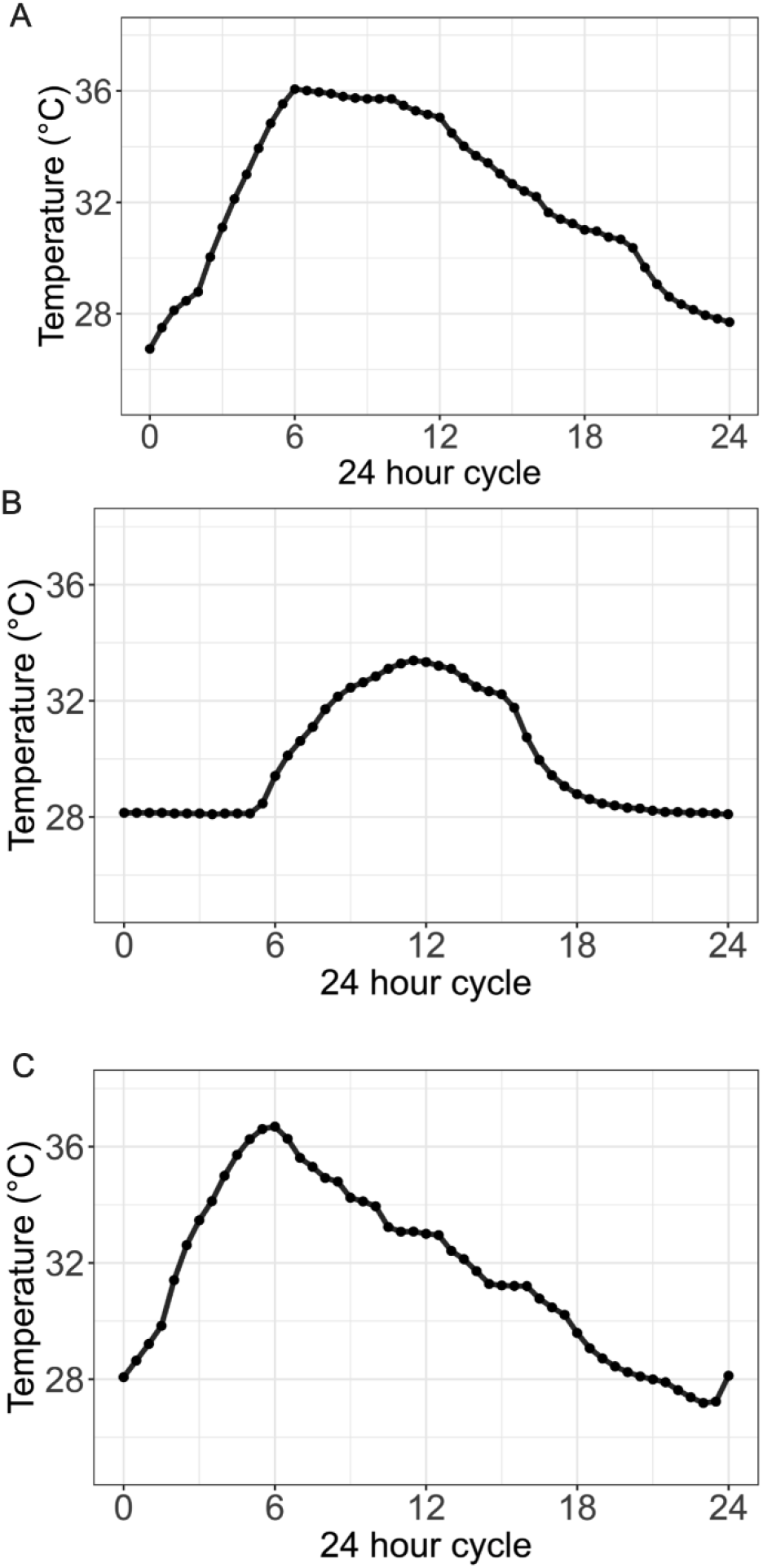
Simulated larval, shaded and semi-shaded temperature cycles. (**A**) A representative 24-hour period of the simulated tropical water-temperature cycle generated from data from water drums known to act as *Ae. aegypti* larvae breeding sites in Trinidad (33). Data collected using a water-proof temperature probe placed in a volume of water equal to that of the larval pans, and left in a dynamic temperature incubator. 24-hour period shaded (**B**) and semi-shaded (**C)** cycles for adult temperatures generated from data collected in urban Kuala Lumpur. Readings are from a temperature probe placed in a dynamic temperature incubator running the replica cycle.

**S2 Figure.**
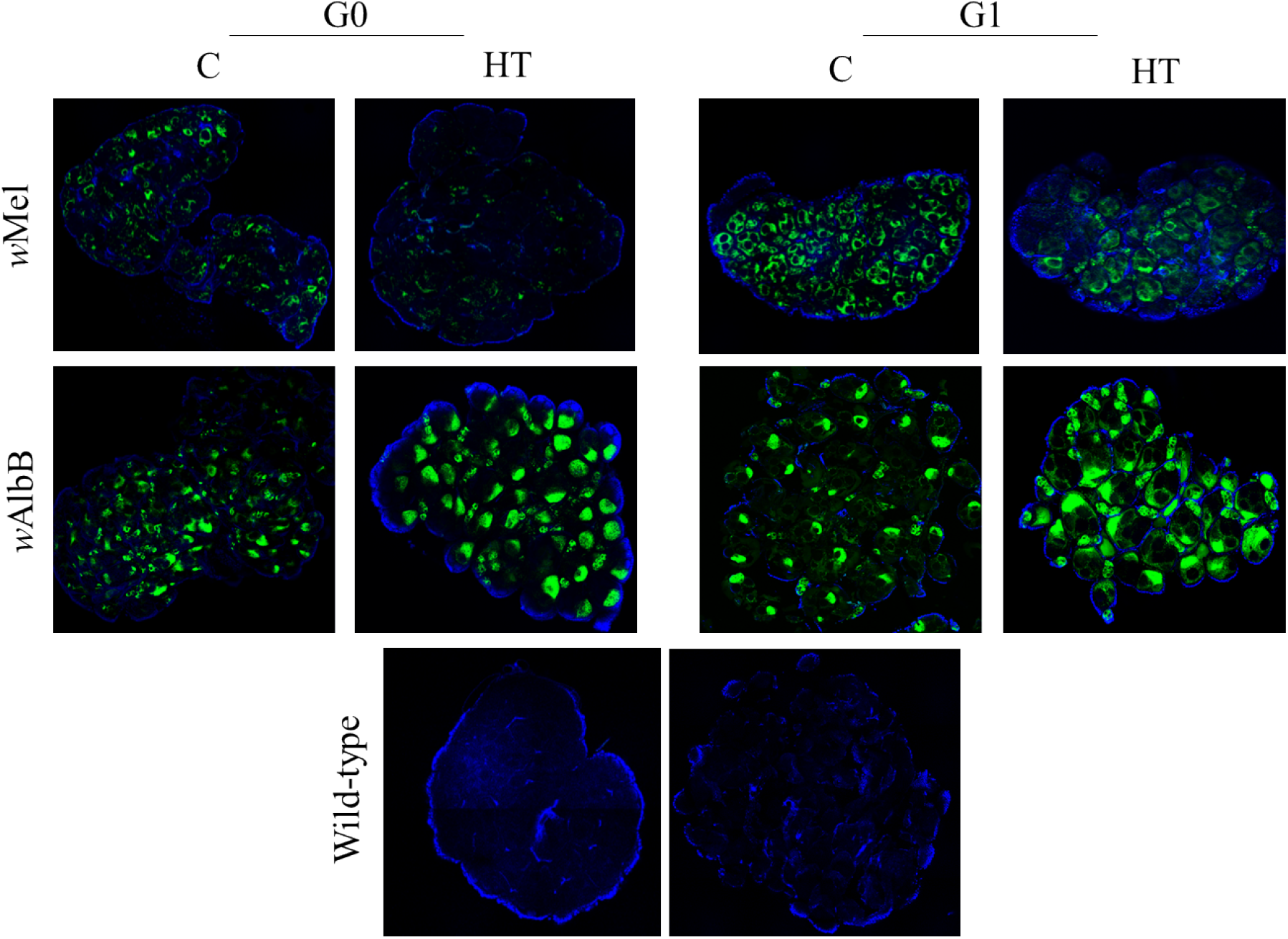
Fluorescent *in situ* hybridization. Visualization of distributions and density reductions of *Wolbachia* (green) in ovaries of 5-days old females from *w*Mel, *w*AlbB and wild-type *Ae. aegypti* females from control and heat-treated groups. Blue stain is DAPI.

**S1 Table:**
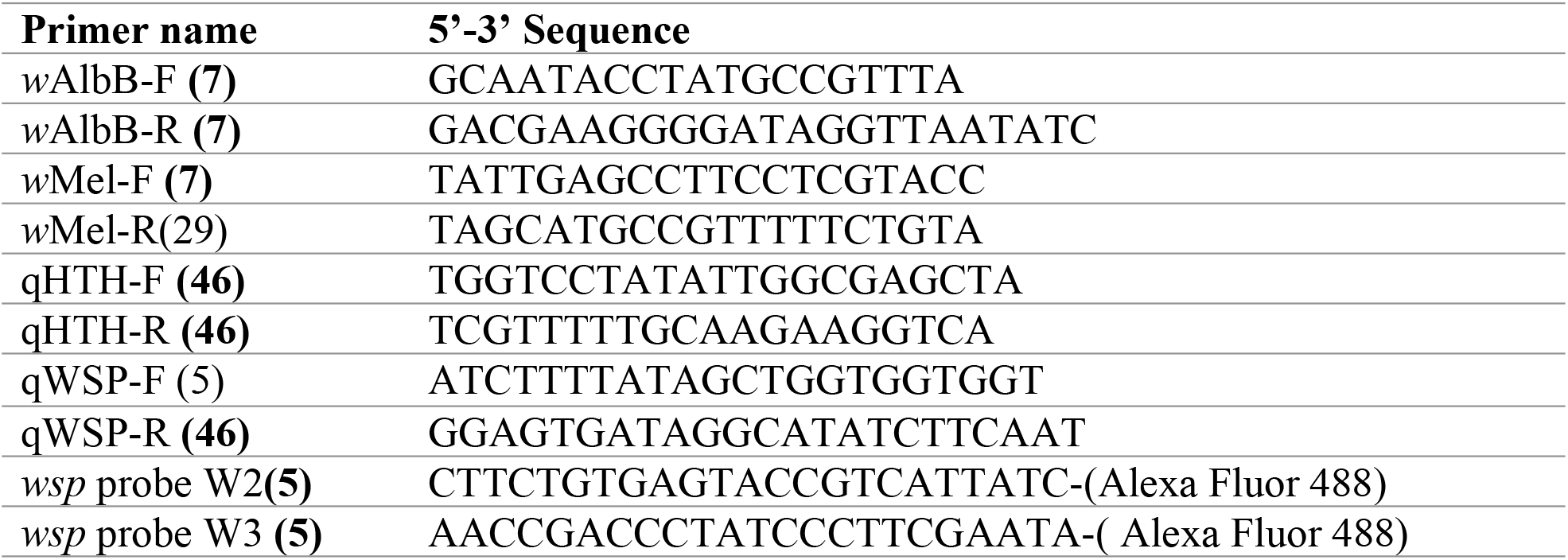
List of sequences of oligonucleotides and probes.

**S2 Table:** Summary of DENV-2 challenge data

|  | Treatment | Tissue | N | Dissemination<br>rate (%) |
| --- | --- | --- | --- | --- |
| **wAlbB** | Heat- treated<br>(HT) | Head and<br>thorax | 24 | 12.5% |
|  |  | Salivary glands | 49 | 8.1% |
|  | Control (C) | Head and<br>thorax | 24 | 0.0% |
| **wMel** | Heat- treated | Head and<br>thorax | 24 | 33.3% |
|  |  | Salivary glands | 50 | 42% |
|  | Control | Head and<br>thorax | 24 | 4.1% |
| **wild-type** | Heat- treated | Head and<br>thorax | 24 | 58.3% |
|  |  | Salivary glands | 46 | 23.9% |
|  | Control | Head and<br>thorax | 24 | 33.3% |

**S3 Table:**
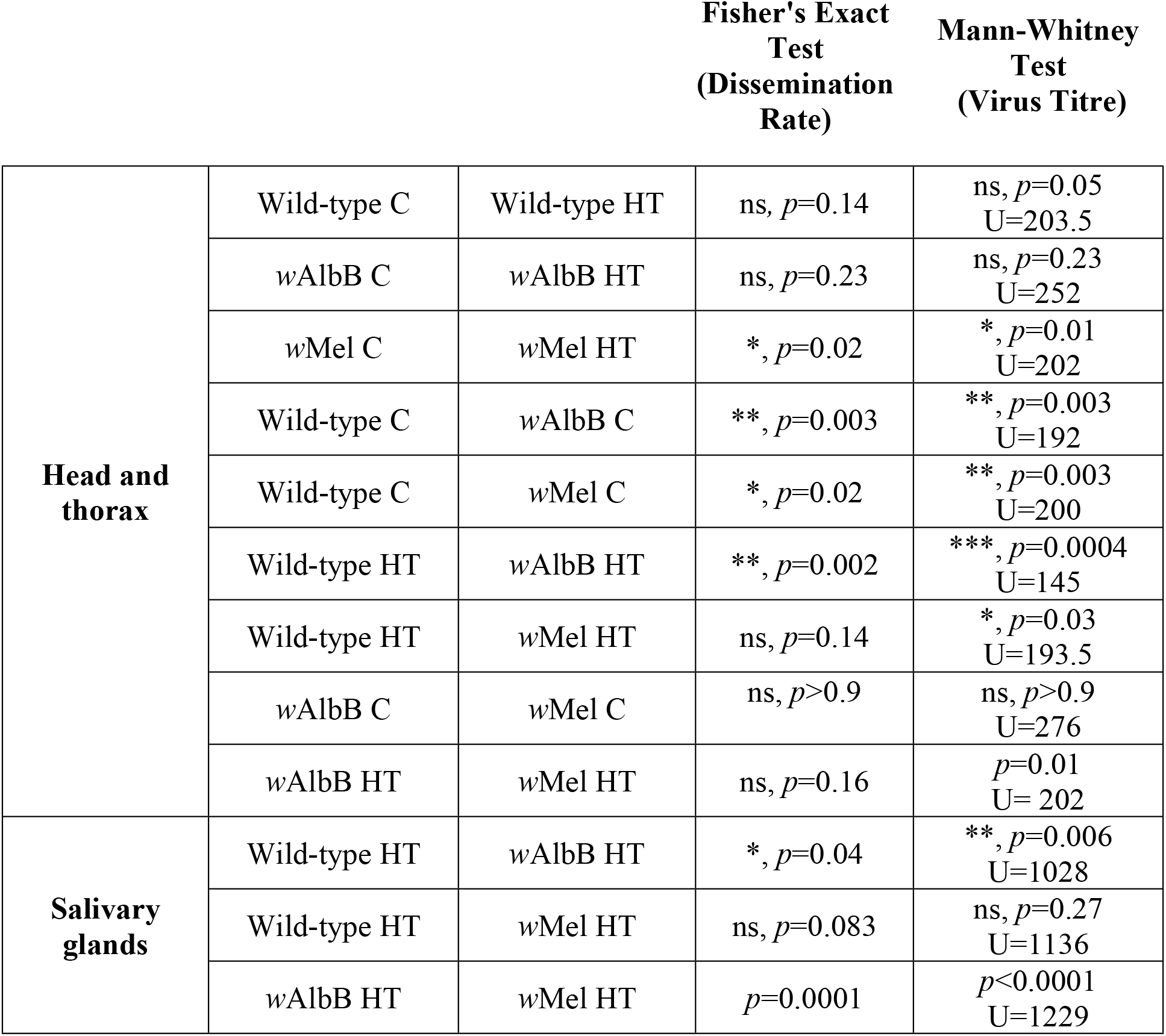
Statistical comparisons between groups and conditions after DENV2 challenge.

## References

1. Ross PA, Callahan AG, Yang Q, Jasper M, Arif MAK, Afizah AN, et al. An elusive endosymbiont: Does Wolbachia occur naturally in Aedes aegypti? Ecol Evol. 2020;10(3):1581–91.

2. Walker T, Johnson PH, Moreira LA, Iturbe-Ormaetxe I, Frentiu FD, McMeniman CJ, et al. The wMel Wolbachia strain blocks dengue and invades caged Aedes aegypti populations. Nature. 2011;476(7361):450–3.

3. Bian G, Xu Y, Lu P, Xie Y, Xi Z. The endosymbiotic bacterium Wolbachia induces resistance to dengue virus in Aedes aegypti. PLoS Pathog. 2010;6(4):e1000833.

4. Hoffmann AA, Ross PA, Rašić G. Wolbachia strains for disease control: ecological and evolutionary considerations. Evol Appl. 2015;8(8):751–68.

5. Moreira LA, Iturbe-Ormaetxe I, Jeffery JA, Lu G, Pyke AT, Hedges LM, et al. A Wolbachia symbiont in Aedes aegypti limits infection with dengue, Chikungunya, and Plasmodium. Cell. 2009;139(7):1268–78.

6. Tan CH, Wong PJ, Li MI, Yang H, Ng LC, O’Neill SL. wMel limits zika and chikungunya virus infection in a Singapore Wolbachia-introgressed Ae. aegypti strain, wMel-Sg. PLoS Negl Trop Dis. 2017;11(5):e0005496.

7. Ant TH, Herd CS, Geoghegan V, Hoffmann AA, Sinkins SP. The Wolbachia strain wAu provides highly efficient virus transmission blocking in Aedes aegypti. PLoS Pathog. 2018;14(1):e1006815.

8. Nazni WA, Hoffmann AA, NoorAfizah A, Cheong YL, Mancini MV, Golding N, et al. Establishment of Wolbachia Strain wAlbB in Malaysian Populations of Aedes aegypti for Dengue Control. Curr Biol. 2019;29(24):4241–8 e5.

9. Ryan PA, Turley AP, Wilson G, Hurst TP, Retzki K, Brown-Kenyon J, et al. Establishment of wMel Wolbachia in Aedes aegypti mosquitoes and reduction of local dengue transmission in Cairns and surrounding locations in northern Queensland, Australia. Gates Open Res. 2019;3:1547.

10. Tantowijoyo W, Andari B, Arguni E, Budiwati N, Nurhayati I, Fitriana I, et al. Stable establishment of wMel Wolbachia in Aedes aegypti populations in Yogyakarta, Indonesia. PLoS Negl Trop Dis. 2020;14(4):e0008157.

11. Joubert DA, Walker T, Carrington LB, De Bruyne JT, Kien DH, Hoang Nle T, et al. Establishment of a Wolbachia Superinfection in Aedes aegypti Mosquitoes as a Potential Approach for Future Resistance Management. PLoS Pathog. 2016;12(2):e1005434.

12. Fraser JE, De Bruyne JT, Iturbe-Ormaetxe I, Stepnell J, Burns RL, Flores HA, et al. Novel Wolbachia-transinfected Aedes aegypti mosquitoes possess diverse fitness and vector competence phenotypes. PLoS Pathog. 2017;13(12):e1006751.

13. Teixeira L, Ferreira A, Ashburner M. The bacterial symbiont Wolbachia induces resistance to RNA viral infections in Drosophila melanogaster. PLoS Biol. 2008;6(12):e2.

14. Hedges LM, Brownlie JC, O’Neill SL, Johnson KN. Wolbachia and virus protection in insects. Science. 2008;322(5902):702.

15. Dutra HL, Rocha MN, Dias FB, Mansur SB, Caragata EP, Moreira LA. Wolbachia Blocks Currently Circulating Zika Virus Isolates in Brazilian Aedes aegypti Mosquitoes. Cell Host Microbe. 2016;19(6):771–4.

16. O’Neill SL, Ryan PA, Turley AP, Wilson G, Retzki K, Iturbe-Ormaetxe I, et al. Scaled deployment of Wolbachia to protect the community from dengue and other Aedes transmitted arboviruses. Gates Open Res. 2018;2:36.

17. Indriani C, Tantowijoyo W, Rancès E, Andari B, Prabowo E, Yusdi D, et al. Reduced dengue incidence following deployments of. Gates Open Res. 2020;4:50.

18. Ekwudu O, Devine GJ, Aaskov JG, Frentiu FD. Wolbachia strain wAlbB blocks replication of flaviviruses and alphaviruses in mosquito cell culture. Parasit Vectors. 2020;13(1):54.

19. Martinez J, Longdon B, Bauer S, Chan YS, Miller WJ, Bourtzis K, et al. Symbionts commonly provide broad spectrum resistance to viruses in insects: a comparative analysis of Wolbachia strains. PLoS Pathog. 2014;10(9):e1004369.

20. Lu P, Bian G, Pan X, Xi Z. Wolbachia induces density-dependent inhibition to dengue virus in mosquito cells. PLoS Negl Trop Dis. 2012;6(7):e1754.

21. McMeniman CJ, Lane RV, Cass BN, Fong AW, Sidhu M, Wang YF, et al. Stable introduction of a life-shortening Wolbachia infection into the mosquito Aedes aegypti. Science. 2009;323(5910):141–4.

22. Min KT, Benzer S. Wolbachia, normally a symbiont of Drosophila, can be virulent, causing degeneration and early death. Proc Natl Acad Sci U S A. 1997;94(20):10792–6.

23. Fraser JE, O’Donnell TB, Duyvestyn JM, O’Neill SL, Simmons CP, Flores HA. Novel phenotype of Wolbachia strain wPip in Aedes aegypti challenges assumptions on mechanisms of Wolbachia-mediated dengue virus inhibition. PLoS Pathog. 2020;16(7):e1008410.

24. Axford JK, Ross PA, Yeap HL, Callahan AG, Hoffmann AA. Fitness of wAlbB Wolbachia Infection in Aedes aegypti: Parameter Estimates in an Outcrossed Background and Potential for Population Invasion. Am J Trop Med Hyg. 2016;94(3):507–16.

25. Hancock PA, White VL, Ritchie SA, Hoffmann AA, Godfray HC. Predicting Wolbachia invasion dynamics in Aedes aegypti populations using models of density-dependent demographic traits. BMC Biol. 2016;14(1):96.

26. Barton NH, Turelli M. Spatial waves of advance with bistable dynamics: cytoplasmic and genetic analogues of Allee effects. Am Nat. 2011;178(3):E48–75.

27. Nguyen TH, Nguyen HL, Nguyen TY, Vu SN, Tran ND, L. TN, et al. Field evaluation of the establishment potential of wMelPop Wolbachia in Australia and Vietnam for dengue control. Parasit Vectors. 2015;8:563.

28. Wernegreen JJ. Mutualism meltdown in insects: bacteria constrain thermal adaptation. Curr Opin Microbiol. 2012;15(3):255–62.

29. Sazama EJ, Ouellette SP, Wesner JS. Bacterial Endosymbionts Are Common Among, but not Necessarily Within, Insect Species. Environ Entomol. 2019;48(1):127–33.

30. Sumi T, Miura K, Miyatake T. Wolbachia density changes seasonally amongst populations of the pale grass blue butterfly, Zizeeria maha (Lepidoptera: Lycaenidae). PLoS One. 2017;12(4):e0175373.

31. Ross PA, Wiwatanaratanabutr I, Axford JK, White VL, Endersby-Harshman NM, Hoffmann AA. Wolbachia Infections in Aedes aegypti Differ Markedly in Their Response to Cyclical Heat Stress. PLoS Pathog. 2017;13(1):e1006006.

32. Ulrich JN, Beier JC, Devine GJ, Hugo LE. Heat Sensitivity of wMel Wolbachia during Aedes aegypti Development. PLoS Negl Trop Dis. 2016;10(7):e0004873.

33. Hemme RR, Tank JL, Chadee DD, Severson DW. Environmental conditions in water storage drums and influences on Aedes aegypti in Trinidad, West Indies. Acta Trop. 2009;112(1):59–66.

34. Ross PA, Hoffmann AA. Continued Susceptibility of the wMel Wolbachia Infection in Aedes aegypti to Heat Stress Following Field Deployment and Selection. Insects. 2018;9(3).

35. K. P. Paaijmans AFGJ, W. Takken, B. G. Heusinkveld, A. K. Githeko, M. Dicke and A. A. M. Holtslag. Observations and model estimates of diurnal water temperature dynamics in mosquito breeding sites in western Kenya. Hydr Process; 2008:22(4789–801).

36. Nainu F, Trenerry A, Johnson KN. Wolbachia-mediated antiviral protection is cell-autonomous. J Gen Virol. 2019;100(11):1587–92.

37. Cattarino L, Rodriguez-Barraquer I, Imai N, Cummings DAT, Ferguson NM. Mapping global variation in dengue transmission intensity. Sci Transl Med. 2020;12(528).

38. Geoghegan V, Stainton K, Rainey SM, Ant TH, Dowle AA, Larson T, et al. Perturbed cholesterol and vesicular trafficking associated with dengue blocking in Wolbachia-infected Aedes aegypti cells. Nat Commun. 2017;8(1):526.

39. Schmidt TL, Barton NH, Rašić G, Turley AP, Montgomery BL, Iturbe-Ormaetxe I, et al. Local introduction and heterogeneous spatial spread of dengue-suppressing Wolbachia through an urban population of Aedes aegypti. PLoS Biol. 2017;15(5):e2001894.

40. Garcia GA, Sylvestre G, Aguiar R, da Costa GB, Martins AJ, Lima JBP, et al. Matching the genetics of released and local Aedes aegypti populations is critical to assure Wolbachia invasion. PLoS Negl Trop Dis. 2019;13(1):e0007023.

41. Carrington LB, Tran BCN, L. NTH, Luong TTH, Nguyen TT, Nguyen PT, et al. Field-and clinically derived estimates of Wolbachia-mediated blocking of dengue virus transmission potential in Aedes aegypti mosquitoes. Proc Natl Acad Sci U S A. 2018;115(2):361–6.

42. Ye YH, Carrasco AM, Dong Y, Sgrò CM, McGraw EA. The Effect of Temperature on Wolbachia-Mediated Dengue Virus Blocking in Aedes aegypti. Am J Trop Med Hyg. 2016;94(4):812–9.

43. Ross PA, Axford JK, Yang Q, Staunton KM, Ritchie SA, Richardson KM, et al. Heatwaves cause fluctuations in wMel Wolbachia densities and frequencies in Aedes aegypti. PLoS Negl Trop Dis. 2020;14(1):e0007958.

44. Ewa Chrostek NM, Marta S Marialva, Luis Teixeira. Wolbachia-conferred antiviral protection is determined by developmental temperature. bioRxiv 2020.06.24.169169: bioRxiv; 2020.

45. Reinhold JM, Lazzari CR, Lahondère C. Effects of the Environmental Temperature on Aedes aegypti and Aedes albopictus mosquitoes: a review. Insects. 2018;9(4).

46. Braig HR, Zhou W, Dobson SL, O’Neill SL. Cloning and characterization of a gene encoding the major surface protein of the bacterial endosymbiont Wolbachia pipientis. J Bacteriol. 1998;180(9):2373–8.

